# Dopaminergic Modulation of Mushroom Body Output Neurons Mediates Nociception-Induced Escape in *Drosophila*

**DOI:** 10.64898/2026.01.04.697536

**Authors:** Chi-Lien Yang, Chia-Wen Chen, Kuan-Lin Feng, Hsiao-Chien Peng, Ming-Chin Wu, Ching-Che Charng, Li-An Chu, Yeong-Ray Wen, Ann-Shyn Chiang

## Abstract

In *Drosophila*, noxious heat is detected by peripheral nociceptors expressing transient receptor potential (TRP) channels, including Painless and TrpA1, and rapidly triggers escape behavior. Although peripheral transduction has been defined in detail, the central circuits and neuromodulatory mechanisms that translate nociceptor activity into escape decisions remain poorly understood. Here, we combine targeted behavioral perturbations with anatomical tracing to delineate a nociception-to-escape pathway that engages dopaminergic modulation of mushroom body (MB) output. Kir2.1-mediated silencing across candidate neurotransmitter systems revealed a specific requirement for MB-innervating dopaminergic neurons (DANs)—particularly subsets within the protocerebral posterior lateral 1 (PPL1) and protocerebral anterior medial (PAM) clusters—for robust nociception-induced escape. Anterograde *trans*-Tango tracing from *painless*- and *trpA1*-expressing nociceptors labeled these MB dopaminergic neurons as direct postsynaptic partners, consistent with convergence of distinct nociceptor inputs onto a shared dopaminergic pathway. Finally, silencing a subset of mushroom body output neurons (MBONs) delayed escape without overtly disrupting baseline locomotion, supporting a model in which dopaminergic signaling recruits MB output to shape defensive action selection. Together, our results define a multi-layer circuit motif linking peripheral nociception to MB-dependent escape and provide a framework for dissecting how neuromodulation gates rapid defensive behaviors.

## Introduction

Nociception relies on specialized sensory neurons (nociceptors) that detect potentially tissue-damaging stimuli, including noxious heat, cold, mechanical forces, and chemical irritants. Transient receptor potential (TRP) channels constitute an evolutionarily conserved family of ion channels that contribute to sensing these modalities across species. Whereas mammalian thermal nociception prominently involves transient receptor potential vanilloid (TRPV) channels (Caterina, et al., 1999; Caterina et al., 1997; Guler et al., 2002; Peier et al., 2002), *Drosophila* primarily uses members of the transient receptor potential ankyrin (TRPA) subfamily to detect noxious heat, notably *TrpA1* and Painless (Hamada et al., 2008; Neely et al., 2011; Sokabe and Tominaga, 2009; Sokabe, Tsujiuchi et al., 2008; Tracey et al., 2003; Zhong et al., 2012). *TrpA1* is activated at warm temperatures and contributes to thermal avoidance across a broad range, whereas Painless exhibits a higher activation threshold and is required for responses to more intense heat (Hamada et al., 2008; Sokabe et al., 2008; Zhong et al., 2012). Consistent with these roles, mutations in *TrpA1* or *painless* impair heat nociception and disrupt escape behaviors in both larvae and adults (Neely et al., 2011; Tracey et al., 2003).

Beyond peripheral sensory organs, drivers based on *painless* and *TrpA1* regulatory sequences label broad neuronal populations in the adult central nervous system. *painless-Gal4* labels neurons innervating the mushroom body (MB), antennal lobe (AL), ellipsoid body, pars intercerebralis, ventral nerve cord, and multiple peripheral structures (Sakai, et al., 2012; Sato et al., 2014; Sun et al., 2009; Wang et al., 2011; Xu et al., 2006), and has been implicated in modulating olfactory processing, sexual behavior, and courtship memory (Sakai et al., 2012; Sato et al., 2014; Wang et al., 2011). *TrpA1-Gal4* labels neurons projecting to the subesophageal zone, near the antennal lobe (including anterior cell thermosensory neurons), superior dorsofrontal protocerebrum, fan-shaped body, and ventral nerve cord, and contributes to temperature preference and circadian locomotor activity (Hamada et al., 2008; Kim et al., 2010; Lee, 2013; Shih and Chiang, 2011). Notably, *painless* and *TrpA1* expression does not fully overlap in peripheral tissues (Mandel et al., 2018), raising the possibility that nociceptive information from these channels converges at downstream nodes in the central brain.

To identify candidate downstream circuits for heat nociception-induced escape, we combined a quantitative laser-based behavioral assay with circuit mapping. We first used a targeted Kir2.1 silencing screen across neurotransmitter-defined populations to identify modulatory systems required for escape. We then applied the trans-Tango transsynaptic labeling system (Talay et al., 2017) to map direct postsynaptic partners of *painless-Gal4* and *TrpA1-Gal4* neurons. These experiments implicated dopaminergic neurons (DANs) as a common convergence point and suggested downstream engagement of MB output pathways. Finally, a focused screen of *split-GAL4* lines targeting defined dopaminergic subsets and MB output neurons (MBONs) identified specific nodes required for nociception-induced escape. Together, our results support a circuit model in which nociceptor-driven signals recruit dopaminergic modulation and MBON output to shape escape behavior.

## Results

### Automated laser-based nociception assay

To quantify nociception-induced escape behavior, we established a laser-based avoidance assay using the Automated Laser Tracking and Optogenetic Manipulation System (ALTOMS) (Wu et al., 2014). To assess nociceptive responses, we tracked individual adult male flies in an acrylic arena while delivering a continuous-wave laser stimulus (473 nm, 10 mW; spot diameter ∼1200 um) to the ventral thorax (Figure 1A). This noxious stimulus typically elicited rapid escape jumps (Figure 1B). We quantified the response by measuring the latency to the first jump within a 180 s stimulation window, while baseline locomotion was assessed via walking velocity. This assay robustly distinguished wild-type *w^1118^* flies from nociception-defective mutants, *painless^1^* and *TrpA1^1^*, which exhibited prolonged jumping latencies (Figure 1C). However, *TrpA1^1^* mutants displayed reduced walking velocity relative to controls (Figure 1D), highlighting the importance of monitoring locomotor confounds when interpreting escape latency.

**Figure 1.**
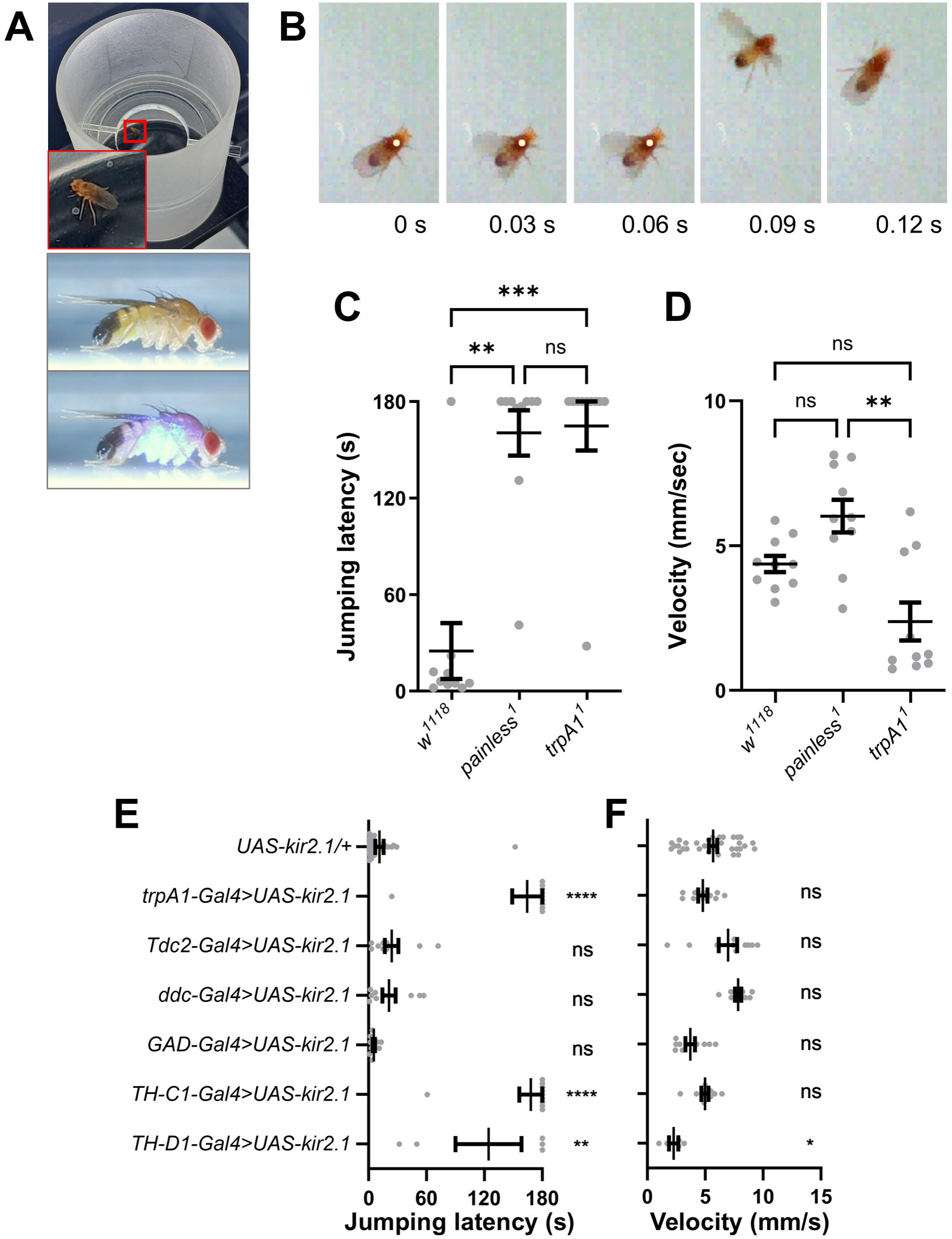
Laser stimulation induces nociceptive escape responses in the male *Drosophila*. (A) In an acrylic arena covered with glass, a freely moving male fly is tracked and targeted by a 473 nm laser beam (10 mW) on the ventral thorax. (B) Representative frames showing a jumping escape response during laser irradiation. (C) Compared with wild-type *w^1118^* controls, *painless^1^* and *trpA1^1^* mutants exhibit prolonged jumping latency during laser irradiation. (D) *trpA1^1^* mutants show reduced locomotor velocity compared with *w^1118^* and *painless^1^*. (E-F) A behavioral screen was performed using *UAS-Kir2.1* to silence candidate *Gal4* drivers targeting distinct neurotransmitter systems. Silencing *trpA1-Gal4* neurons served as a positive control and increased jumping latency. The *painless-Gal4* driver was excluded because the *Gal4* insertion disrupts endogenous *painless* function. Silencing DANs labeled by *TH-C1-Gal4* or *TH-D1-Gal4* increased jumping latency (E); *TH-D1-Gal4* also reduced walking velocity (F). Data were analyzed by Kruskal-Wallis test. Significance: ns, not significant; *P < 0.05, **P < 0.01, ***P < 0.001, ****P < 0.0001.

### Dopaminergic neurons are required for nociception-induced escape behavior

To identify neuronal populations that modulate nociception-induced escape, we chronically silenced candidate *Gal4*-defined neurons by expressing the inward-rectifying potassium channel Kir2.1. As a positive control, silencing *trpA1-Gal4* neurons significantly increased jumping latency without affecting escape velocity (Figure 1E, F). We next performed a neurotransmitter-specific screen by silencing octopaminergic, serotonergic, GABAergic, and dopaminergic neurons (DANs). Silencing GABAergic (*GAD-Gal4*), octopaminergic (*Tdc2-Gal4*), or mixed dopaminergic/serotonergic neurons (*ddc-Gal4*) did not significantly alter either jumping latency or velocity. In contrast, silencing DANs using *TH-C1-Gal4* or *TH-D1-Gal4* markedly increased jumping latency (Figure 1E). While *TH-C1-Gal4* silencing had no significant effect on velocity, *TH-D1-Gal4* silencing reduced locomotion velocity (Figure 1F), indicating that distinct subsets of DANs contribute to escape initiation and partially overlap with locomotor control circuits. Collectively, these findings identify specific dopaminergic neuron subsets as key modulators of nociception-induced escape behavior.

### Dopaminergic neurons are direct downstream targets of *painless-* and *trpA1-*expressing neurons

To examine connectivity between heat nociceptor populations and dopaminergic circuits, we mapped downstream targets of *painless-Gal4* and *trpA1-Gal4* neurons using *trans*-Tango (Talay et al., 2017) combined with tyrosine hydroxylase (TH) immunostaining (Figure 2A, B). Both tracing experiments revealed *trans*-Tango signal colocalized with multiple TH-positive clusters, indicating that DANs are among the direct postsynaptic partners of both *painless*- and *trpA1*-expressing neurons. *Trans*-Tango labeling was observed in protocerebral anterior lateral (PAL) cluster, protocerebral anterior medial (PAM) cluster, protocerebral posterior lateral clusters (including PPL1, PPL2ab, PPL2c), protocerebral posterior medial clusters (including PPM1, PPM2, PPM3) (Figure 2A, B), several of which innervate the mushroom body neuropil (Mao and Davis, 2009). These anatomical data support the idea that distinct nociceptor populations can converge on dopaminergic pathways that are positioned to influence mushroom body processing.

**Figure 2.**
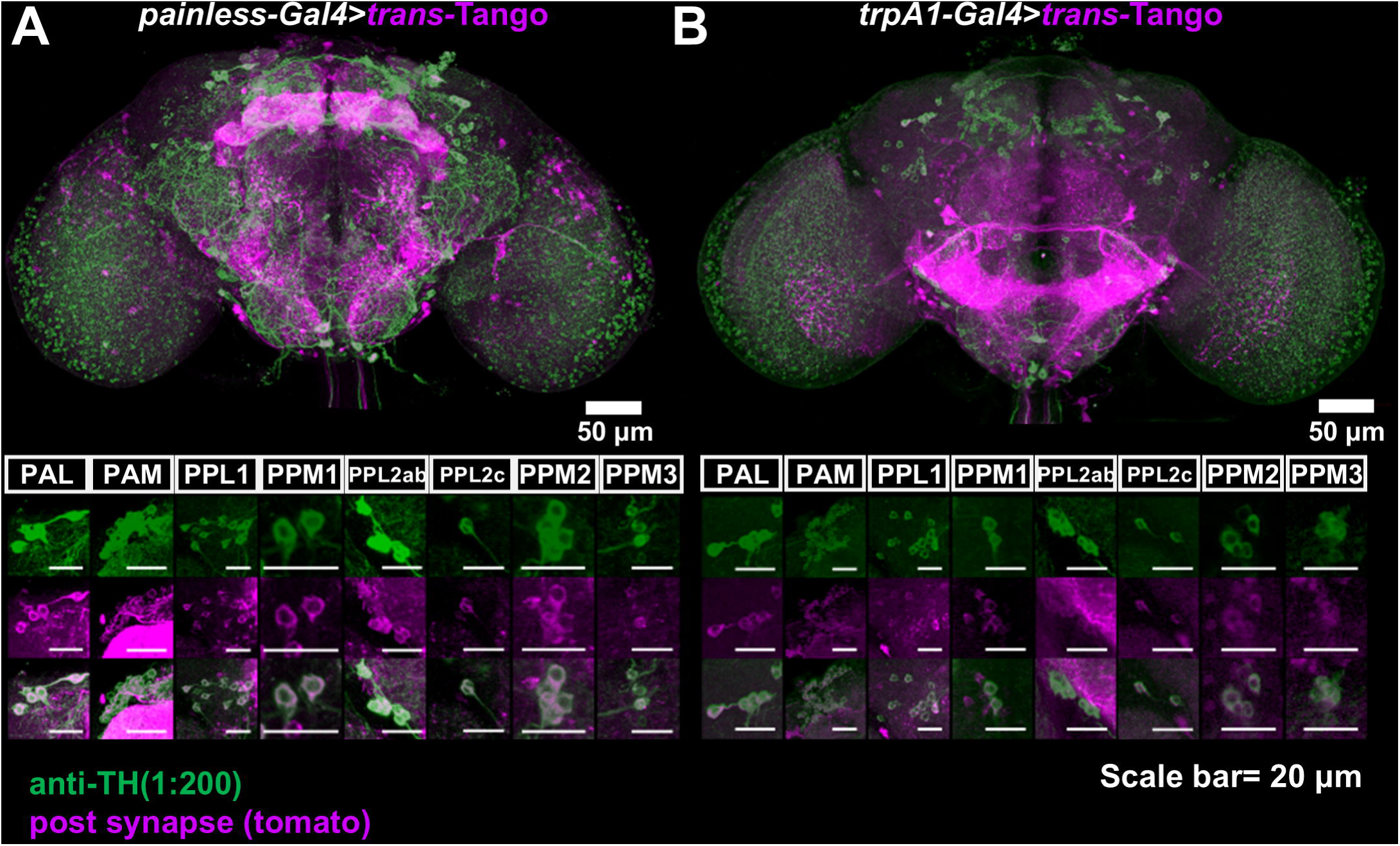
Anatomical tracing implicate dopaminergic neurons in nociception-induced escape. (A-B) *trans*-Tango labeling reveals direct downstream partners of nociceptor-associated neurons. Downstream targets of (A) *painless-Gal4* and (B) *trpA1-Gal4* neurons were labeled by *trans*-Tango (magenta), and DANs were visualized by TH immunostaining (green). Colocalization indicates that multiple dopaminergic clusters (PAL, PAM, PPL1, PPM1, PPL2ab, PPL2c, PPM2, and PPM3) are labeled as direct postsynaptic targets.

### Specific PPL1 and PAM dopaminergic subsets are essential for nociception-induced escape behavior

Because MB-innervating PPL1 and PAM DANs have been implicated in aversive reinforcement and action selection (Aso et al., 2010; Masek et al., 2015), we next screened *split-GAL4* lines that target defined PPL1 or PAM subsets (Aso et al., 2014). Silencing PPL1-targeting lines MB060B, MB065B, MB296B, MB304B, and MB308B increased jumping latency while largely preserving walking velocity (Figure 3A,B). Silencing MB058B also increased latency but was accompanied by reduced velocity, suggesting broader effects on locomotion. Within the PAM cluster, silencing MB213B or MB301B increased jumping latency without significantly affecting velocity (Figure 3A,B). Notably, silencing other MB-innervating dopaminergic subsets did not affect jump latency, indicating functional heterogeneity within both clusters. Together, these results show that multiple—though not all—PPL1 and PAM dopaminergic subtypes are required for timely nociception-induced escape.

**Figure 3.**
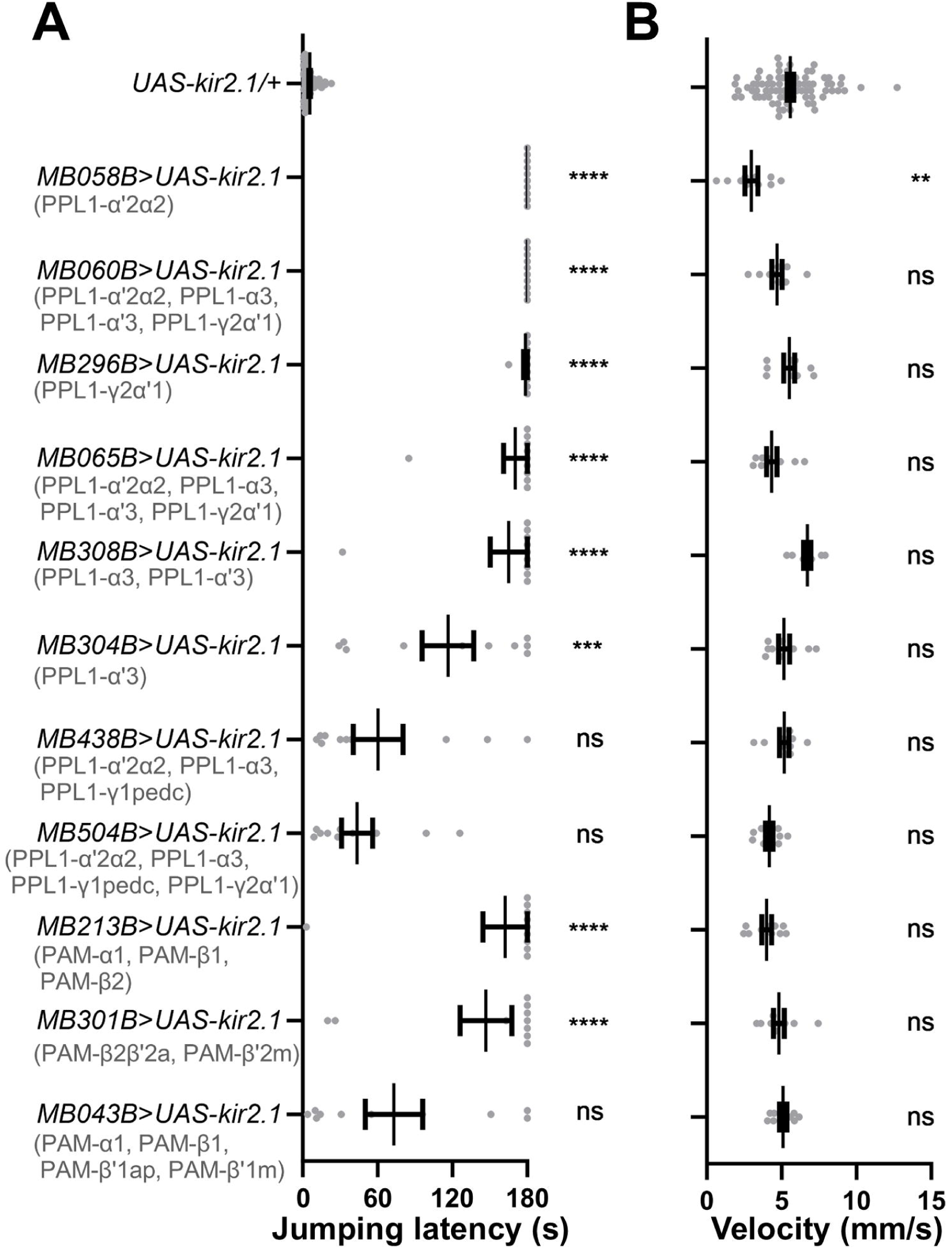
Specific subsets of PPL1 and PAM dopaminergic neurons are required for nociception-induced escape. Targeted silencing of PPL1 subsets using *split-GAL4* drivers MB058B, MB060B, MB065B, MB296B, MB304B, and MB308B increased jumping latency. Silencing PAM subsets using MB213B and MB301B similarly increased jumping latency. Quantification of (A) jumping latency and (B) velocity is shown. Statistics: Kruskal-Wallis test. Significance: ns, not significant; *P < 0.05, **P < 0.01, ***P < 0.001, ****P < 0.0001.

### Mushroom body output neurons are required for nociception-induced escape behavior

Because PPL1 and PAM DANs densely innervate MB lobes in a sector-specific manner, we next asked whether MB output neurons (MBONs) innervated distinct sectors participate in nociception-evoked escape. We expressed *UAS-Kir2.1* in a panel of MBON *split-GAL4* lines (Aso et al., 2014) and quantified escape latency and walking velocity. Silencing multiple MBON classes increased jump latency to varying extents, generally without a corresponding reduction in baseline velocity (Figure 4A, B), arguing against a nonspecific locomotor impairment. The most pronounced effects were observed for MB082C, MB083C, MB298B, MB433B, and MB434B. Together, these results indicate that distinct MB output channels are differentially required for timely nociception-induced escape.

**Figure 4.**
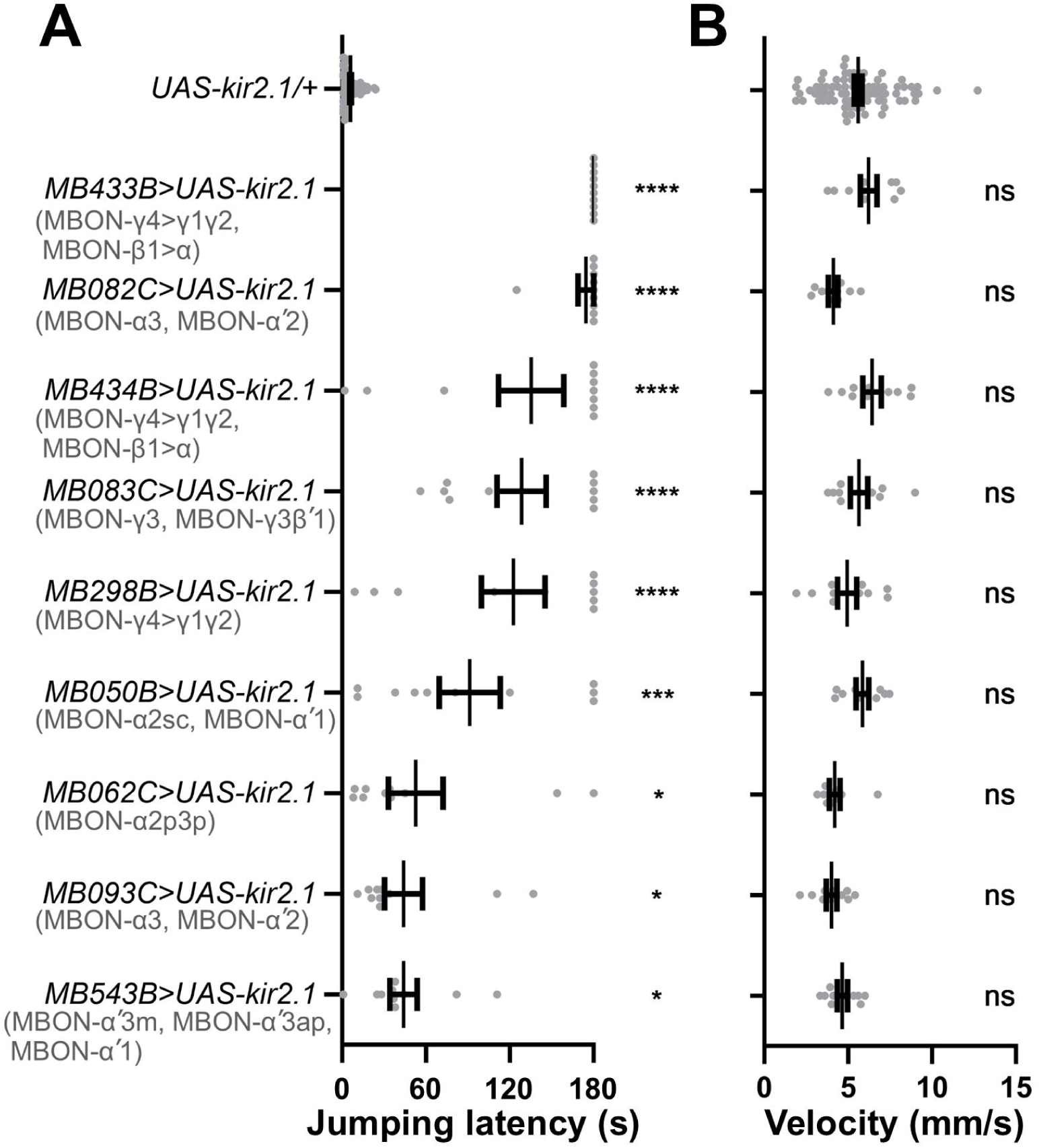
Mushroom body output neurons (MBONs) contribute to nociception-induced escape. Silencing MBONs increased jumping latency, with the largest effects observed for MB082C, MB083C, MB298B, MB433B, and MB434B. Statistics: jumping latency was analyzed by Kruskal-Wallis test; velocity was analyzed by one-way ANOVA where indicated. Significance: ns, not significant; *P < 0.05, **P < 0.01, ***P < 0.001, ****P < 0.0001.

### A circuit model for nociception-induced escape

Integrating our behavioral and anatomical results, we propose a multi-layer circuit model for nociception-induced escape (Figure 5). In this model, acute noxious thermal stimulation is detected by peripheral nociceptors expressing *painless* or *trpA1*. These signals are relayed to central circuits that recruit DANs; *trans*-Tango tracing places multiple dopaminergic clusters downstream of both nociceptor populations. Consistent with this anatomical organization, silencing defined PPL1 and PAM dopaminergic subsets impair timely escape, and silencing MB output neurons (MBONs) associated with MB compartments receiving these dopaminergic inputs similarly delays escape. To systematically evaluate the functional weight of these microcircuits, we calculated a ‘Behavioral Potency Level’ for each compartment by averaging the behavioral responses across all relevant drivers. This analysis stratified MB compartments into high, moderate, and low impact tiers, revealing a modular logic underlying the escape response. For dopaminergic modulation, PPL1-mediated inputs to the α’1, α’3 and γ2 compartments exhibited the highest potency, whereas the β2 and β’2 compartments emerged as the critical targets for PAM neurons. Downstream of these inputs, the output pathways were dominated by MBONs projecting from β1 (to the α lobe) and the recurrent γ4 (to γ1/γ2) circuit. This input-output mapping identifies β1 and γ2 as critical integration hubs.To validate the structural basis of these functional interactions, we analyzed the direct synaptic connectivity between DANs and MBONs using the NeuPrint male CNS connectome (Sturner et al., 2025). We found that synaptic strengths varies distinctively across these compartments. The β1 and β’1 compartments exhibit high connectivity weights (> 1000 synapses), mirroring their strong functional output. In contrast, compartments in the α lobe show moderate connectivity (< 1000 synapses). Intriguingly, despite the high functional impact of γ2, the γ1 and γ2 compartments display low direct synaptic connectivity (< 100 synapses). This structural sparsity suggests that the profound modulatory effect of PPL1 on γ2 may rely on volume transmission or highly effective sparse synapses, rather than dense wired connectivity.

**Figure 5.**
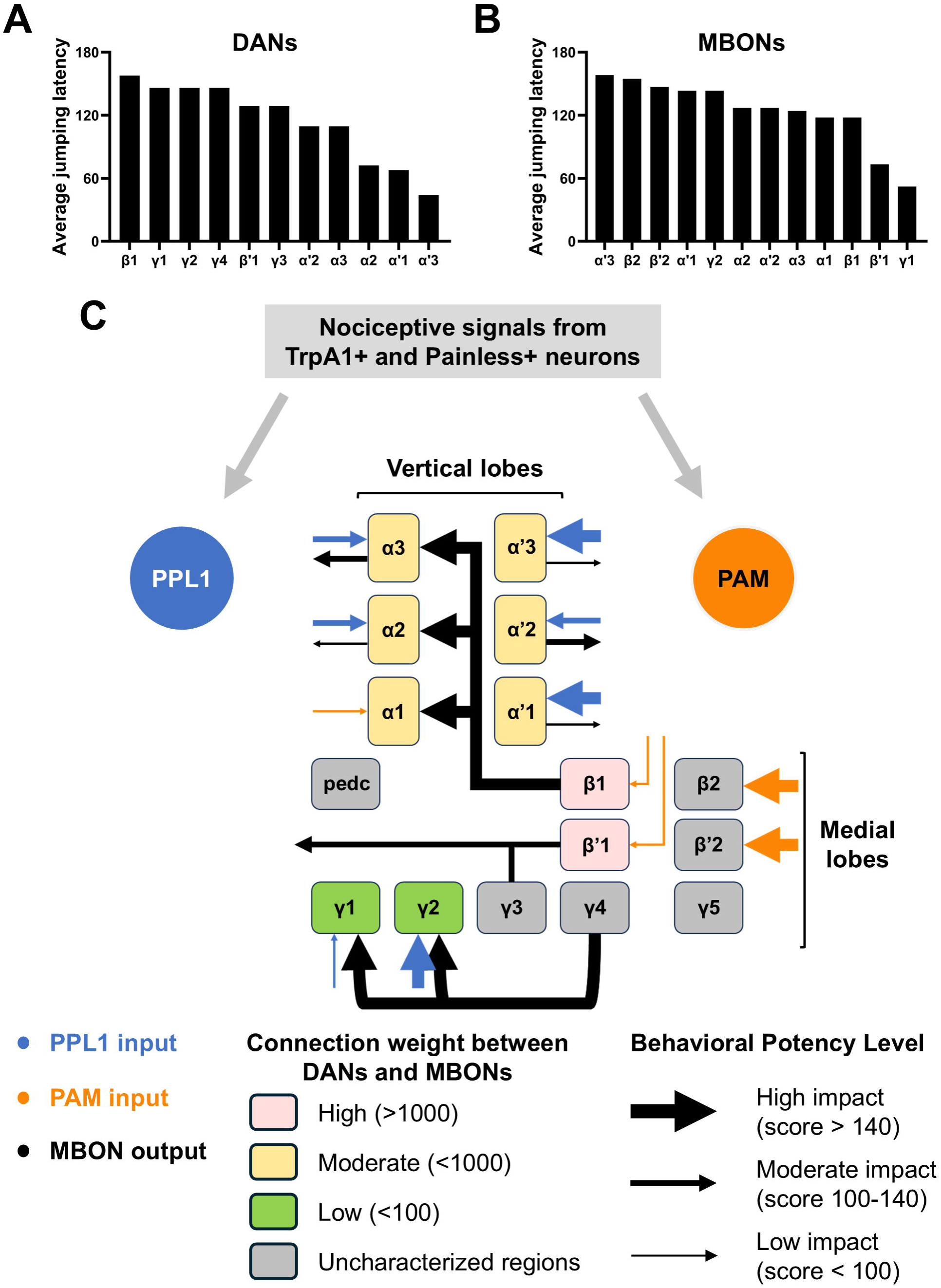
Working model of a MB–centered circuit for nociception-induced escape jumping. Noxious stimuli detected through *painless**-*** and *trpA1*-dependent sensory pathways are proposed to recruit defined subsets of DANs labeled by *TH-C1-GAL4*, *TH-D1-GAL4***, a**nd MB compartment-specific *split-GAL4* drivers. These DANs (PPL1, blue; PAM, orange) innervate discrete MB compartments and are hypothesized to modulate compartmental MB computations to shape the output of MB output neurons (MBONs; black) that control escape-jump latency. This schematic integrates (i) behavioral silencing screens identify compartments required for normal escape performance and (ii) *trans*-Tango anterograde transsynaptic mapping was used to nominate downstream MBON connectivity. Arrow thickness represents Behavioral Potency Level (behavioral impact), defined as the mean escape-jump latency obtained across all driver lines targeting a given compartment, binned into high, moderate, or low effect tiers. Among DAN inputs, PPL1 innervation of α′1, α′3, and γ2 shows the largest behavioral impact, with additional contributions from α2, α3, and α′2 compartments. Within the PAM cluster, inputs to β2 and β′2 are most strongly associated with escape performance. For MBON outputs, inter-compartment MBONs projecting from β1 to the α lobe and from γ4 to γ1/γ2 exert the strongest effects, while outputs from β′1 and γ3 also contribute. Together, the convergence of dopaminergic drive and MBON routing nominates β1 and γ2 as candidate integration hubs: β1 couples strong PAM input with output to the α lobe, potentially reconfiguring downstream action selection, whereas γ2 acts as a convergence zone receiving PPL1 modulation and recurrent MBON-driven signals and is indispensable for the escape response.

## Discussion

By combining a quantitative escape assay with cell-type–targeted silencing and transsynaptic anatomical tracing, we identify dopaminergic modulation of MB compartments and their output pathways as central components of nociception-induced escape in adult *Drosophila*. Our results support a multilayer circuit architecture in which nociceptor-associated pathways recruit discrete subsets of dopaminergic PPL1 and PAM neurons, which in turn bias MB compartmental processing to shape MB output neuron (MBON) activity and ultimately determine the timing of escape. This framework is consistent with extensive evidence that MB compartments are functionally specialized and that MBON ensembles can gate action selection and valence-dependent behavior (Aso et al., 2014).

The involvement of the MB further implies that nociceptive escape is not exclusively a hard-wired reflex, but can be centrally integrated and state-dependently gated. The MB is well positioned to combine sensory evidence with internal-state variables (e.g., hunger, arousal, sleep pressure) and prior experience to tune behavioral thresholds and decision policies (Sitaraman et al., 2015; Suarez-Grimalt et al., 2024; Tsao et al., 2018; Vanderheyden et al., 2019). In parallel, intense or spatially localized threats may preferentially engage faster routes from brain to ventral nerve cord (VNC) via descending pathways, enabling rapid, stereotyped motor programs when speed is paramount (Medeiros et al., 2024; von Reyn et al., 2017). An important next step will be to determine how stimulus intensity, context, and internal state shift the relative weighting of MB-dependent computation versus more direct VNC-linked escape circuitry.

Distinct dopaminergic circuits are known to encode reinforcement and regulate both learned and innate actions. In canonical olfactory learning, specific PPL1 DANs convey aversive teaching signals, and PAM DANs are classically associated with reward, though subsets can support aversive reinforcement depending on conditions and task structure (Aso et al., 2010; Claridge-Chang et al., 2009). In our laser-based assay, silencing some previously characterized aversive-related DAN subsets (e.g., PPL1-γ1pedc lines) did not affect escape latency, whereas perturbing other PPL1 subsets and PAM-innervating lines produced robust phenotypes. Together, these observations suggest that noxious heat recruits a partially distinct dopaminergic ensemble compared with electric shock or bitter reinforcement, consistent with modality- and context-specific routing of aversive signals within the broader DAN system (Aso et al., 2010; Masek et al., 2015).

Our functional mapping also reveals nontrivial input–output transformations across MB compartments. In β1, dopaminergic input produced relatively weak behavioral effects, whereas silencing the corresponding MBON output strongly impaired escape, pointing to a potent downstream action channel that is not simply predicted by local DAN potency. Conversely, in α′1 and α′3, dopaminergic input was highly potent while the MBON outputs tested showed minimal impact, raising the possibility that these compartments act primarily as gates (e.g., by modulating gain, timing, or competition among MBON pathways) rather than directly driving motor commands, or that our genetic access did not capture the full complement of relevant output channels. These dissociations fit with connectome-level evidence that MBON targets are distributed and that compartmental influence can be expressed through multi-synaptic routing rather than one-to-one DAN→MBON coupling (Li et al., 2020).

Notably, our analysis identifies γ2 as a functional hub. Despite its high behavioral potency, available connectomic reconstructions of the adult MB indicate that some DAN–MBON pairings can be comparatively sparse at the level of direct synaptic contacts, suggesting that circuit impact may arise through indirect pathways and/or neuromodulatory mechanisms not captured by simple “wired connectivity” metrics (Li et al., 2020). One attractive hypothesis is that dopaminergic influence in γ2 may partially rely on extrasynaptic/volume transmission, allowing broad modulation even when anatomically resolved synapses are limited (Liu et al., 2021; Ueno and Kume, 2014). Testing this idea will require experiments that directly relate dopamine dynamics and receptor activation to compartmental physiology during escape.

Finally, while *trans*-Tango provides an efficient entry point to nominate candidate downstream partners of nociceptor-associated neurons, its outputs require careful interpretation. *trans*-Tango is based on a synthetic signaling cascade that reports chemical synaptic connectivity and, by design, does not capture electrical coupling (Talay et al., 2017); reporter sensitivity and expression can also generate false negatives, and anatomical labeling alone cannot establish functional coupling in a specific behavioral context (Talay et al., 2017). For this reason, the functional requirements revealed by Kir2.1-based perturbations are essential for assigning circuit relevance. Going forward, combining acute activation/inhibition, compartment-resolved calcium or voltage imaging, and synapse-level connectomics will be crucial to validate directionality, identify intermediary relays, and define how dopaminergic modulation of specific MB compartments reshapes MBON output to control nociception-induced escape.

## Materials and Methods

### Fly strains and husbandry

Flies were maintained on standard cornmeal medium in plastic vials (2.5 cm diameter × 9.5 cm height) at 25 °C, 70% relative humidity, under a 12 h:12 h light/dark cycle. Unless otherwise indicated, experiments were performed on adult males, 5–10 days post-eclosion.

Stocks were obtained from the Bloomington Drosophila Stock Center (BDSC) unless noted: *w^1118^* (5905), *painless^1^* (27895), *trpA1^1^* (26504), *painless-Gal4* (27894), *trpA1-Gal4* (27593), *Tdc2-Gal4* (9313), *GAD-Gal4* (51630), *ddc-Gal4* (7009), *UAS-GFP* (5137 and 5130), *trans*-Tango (77481), MB043B (68304), MB050B (68365), MB058B (68278), MB060B (68279), MB065B (68281), MB082C (68286), MB083C (68287), MB093C (68289), MB213B (68273), MB296B (68308), MB298B (68309), MB301B (68311), MB304B (68367), MB308B (68312), MB433B (68324), MB434B (68325), MB438B (68326), and MB504B (68329), MB543B (68335). UAS-Kir2.1 (inward-rectifying K^+^ channel used for neuronal silencing) was a gift from Regina Vittore; its use as a chronic silencer in *Drosophila* is well established (Hodge, 2009). *TH-C1-GAL4* and *TH-D1-GAL4* were gifts from Mark Wu. MB062C was obtained from FlyLight (Janelia Research Campus, Ashburn, Virginia, USA).

#### Note for transparency

Because *painless-Gal4* is an insertion in the *painless* locus, it can disrupt endogenous gene function; therefore, it was not used for functional silencing assays (see Figure 1 legend).

### Immunostaining

Adult brains were dissected in phosphate-buffered saline (PBS) and fixed in 4% paraformaldehyde at room temperature for 30 min. Samples were blocked /permeabilized in PBS containing 2% Triton X-100 and 10% normal goat serum. To enhance antibody penetration, brains were degassed under vacuum (-70 mmHg) for 10 min, repeated for four cycles. Brains were incubated at 4 °C for 48 h with primary antibodies diluted in PBS + Triton: (i) mouse 4F3 anti-DLG (1:10, Hybridoma Bank, University of Iowa, Iowa City, IA, USA, Cat# AB_528203) or (ii) mouse anti-TH (1:500; ImmunoStar, Hudson, WI, USA, Cat# 22941). After three washes, samples were incubated with goat anti-mouse Biotin-XX (1:200, Invitrogen, Thermo Fisher Scientific, Waltham, MA, USA, Cat# B-2763) at 4 °C for 24 h, washed, then incubated with Alexa Fluor 635 conjugate –streptavidin (1:500; Thermo Fisher Scientific, Waltham, MA, USA, Cat# S32364) at 4 °C for 24 h. Brains were washed and optically cleared in FocusClear (CelExplorer, HsinChu, Taiwan, Cat# FC-101) prior to mounting between two coverslips separated by ∼250 µm spacers.

### Confocal imaging

Brains were imaged on a Zeiss LSM 710 confocal microscope (Carl Zeiss, Oberkochen, Germany) using a 40× C-Apochromat water-immersion objective (NA 1.2; working distance 220 µm).

### Nociception-induced escape assay

A single male fly was introduced into an acrylic arena (20 mm diameter, 3 mm height) covered with a glass lid. Flies were tracked in real time using ALTOMS (provided by Dr. Ann-Shyn Chiang’s lab), an automated laser-tracking and targeting system, and stimulated with a continuous-wave 473 nm laser (10 mW; spot diameter ∼1200 µm) directed to the ventral thorax for 180 s (Wu et al., 2014). Jumping latency (time to first jump during stimulation) was used as the primary readout of nociception-induced escape. Walking velocity was computed from tracking trajectories to assess locomotor effects.

### Statistical analysis

Jumping latency distributions were analyzed using non-parametric tests (no assumption of normality). For comparisons across multiple genotypes, a Kruskal–Wallis test was used. For pairwise comparisons, two-tailed Mann–Whitney test was performed at a 95% confidence level. Where velocity data met parametric assumptions, one-way ANOVA was used as indicated in figure legends. Test type and significance thresholds are reported in the corresponding figure legends.

## Author contributions

**Chi-Lien Yang:** conceptualization, formal analysis, investigation, methodology, data curation, visualization, writing–original draft, writing–review and editing. **Kuan-Lin Feng:** conceptualization, methodology, writing–review and editing. **Hsiao-Chien Peng:** formal analysis, investigation. **Ming-Chin Wu:** software. **Ching-Che Charng:** formal analysis, data curation. **Li-An Chu:** conceptualization, writing–review and editing. **Chia-Wen Chen:** conceptualization, funding acquisition, writing–review and editing. **Yeong-Ray Wen:** conceptualization, funding acquisition, writing–review and editing. **Ann-Shyn Chiang:** conceptualization, funding acquisition, project administration, resources, supervision, validation, visualization, writing–review and editing.

All authors contributed to and approved the final version of the manuscript.

## Acknowledgments

We thank the Bloomington *Drosophila* Stock Center (supported by NIH P40OD018537), Mark Wu (JHU, Neurology), and FlyLight (Janelia research campus) for providing fly stocks.

## Funding

This work was supported by the National Science and Technology Council, Taiwan (Grants112QR001EK and MOST 105-2314-B-039-006); the Brain Research Center of NTHU under the Featured Areas Research Center Program within the framework of the Higher Education Sprout Project, funded by the Ministry of Education, Taiwan; China Medical University Hospital (Grant DMR-108-213 and CMU106-N-26); and Asia University Hospital (Grant 10651003). Additional support was provided by the Peng Education and Welfare Foundation. The funders had no role in study design, data collection and analysis, decision to publish, or preparation of the manuscript.

## Conflicts of Interest

The authors declare no conflicts of interest.

## Generative AI statement

The authors declare that Generative AI tools, specifically ChatGPT (OpenAI) and Gemini (Google), were used to assist in the preparation of this manuscript. These tools were utilized solely for text paraphrasing and grammar correction to improve clarity and readability. The authors have carefully reviewed and verified all AI-generated content and take full responsibility for the final version of the manuscript.

## Data Availability Statement

The data that support the findings of this study are available from the corresponding author upon reasonable request.

## Notes

### Competing Interest Statement

The authors have declared no competing interest.

### Summary of Updates

Section on Results updated to include full names of dopaminergic neuron subsets; Section on Materials and Methods updated to provide detailed information on antibody and instrument manufacturers; Author list arrangement and affiliations updated.

